# Protein arginine methyltransferase 5 (Prmt5) localizes to chromatin loop anchors and modulates expression of genes at TAD boundaries during early adipogenesis

**DOI:** 10.1101/2023.06.13.544859

**Authors:** Sabriya A. Syed, Kristina Shqillo, Ankita Nand, Ye Zhan, Job Dekker, Anthony N. Imbalzano

## Abstract

Protein arginine methyltransferase 5 (Prmt5) is an essential regulator of embryonic development and adult progenitor cell functions. Prmt5 expression is mis-regulated in many cancers, and the development of Prmt5 inhibitors as cancer therapeutics is an active area of research. Prmt5 functions via effects on gene expression, splicing, DNA repair, and other critical cellular processes. We examined whether Prmt5 functions broadly as a genome-wide regulator of gene transcription and higher-order chromatin interactions during the initial stages of adipogenesis using ChIP-Seq, RNA-seq, and Hi-C using 3T3-L1 cells, a frequently utilized model for adipogenesis. We observed robust genome-wide Prmt5 chromatin-binding at the onset of differentiation. Prmt5 localized to transcriptionally active genomic regions, acting as both a positive and a negative regulator. A subset of Prmt5 binding sites co-localized with mediators of chromatin organization at chromatin loop anchors. *Prmt5* knockdown decreased insulation strength at the boundaries of topologically associating domains (TADs) adjacent to sites with Prmt5 and CTCF co-localization. Genes overlapping such weakened TAD boundaries showed transcriptional dysregulation. This study identifies Prmt5 as a broad regulator of gene expression, including regulation of early adipogenic factors, and reveals an unappreciated requirement for Prmt5 in maintaining strong insulation at TAD boundaries and overall chromatin organization.

## INTRODUCTION

Stem cells are characterized by unique gene expression profiles and the requirement to rapidly undergo transcriptional reprogramming upon appropriate developmental or differentiation signaling. The Prmt5 arginine methyl transferase modulates stem and precursor cell function. Prmt5 plays an important role in maintaining precursor stemness of embryonic and neural stem cells and erythrocyte and hematopoietic progenitor cells(1–4). A potential mechanism by which Prmt5 contributes to stem cell maintenance is by transcriptionally repressing cell cycle inhibitors and promitotic genes(1,2,5). Other studies have suggested a role for Prmt5 as a molecule that facilitates cell differentiation by transcriptional activation through its interactions with co-activators Wdr5(6,7), the mammalian SWI/SNF (mSWI/SNF) chromatin remodeling enzyme(8,9) and the Mediator complex(10) which is a known effector of genome organization(11).

Prmt5 catalyzes symmetric dimethylation of arginine residues on substrate molecules and can regulate transcription through the dimethylation of histone substrates like H4R3 and H3R8(12–14). It also acts on non-histone substrates to control transcription, RNA processing, DNA repair, and protein stability(15). We demonstrated previously that Prmt5 acts as a transcriptional activator during differentiation of cell types derived from mesenchymal stem cells such as myoblasts(16) and pre-adipocytes(8,17). Prmt5 activated gene transcription by facilitating the interaction of the mSWI/SNF chromatin remodeling enzyme with differentiation-specific gene promoters(8,9,16). During adipogenesis, Prmt5 also mediated higher-order chromatin interactions within or between specific loci, including enhancer-promoter looping(^9^), and cis- and trans- chromosomal interactions between actively expressed genes(^18^). The regulation of chromatin looping events involved Prmt5 association with Med1(9), a component of the Mediator complex that has a genome-wide role in enhancer-promoter looping events and potentiating gene transcription(19). During adipogenesis, dynamic and genome-wide changes in promoter-enhancer looping events occur(20). Prior studies have shown that adipogenic loci *Pparγ2*, *Klf4*, *Id2*, *and Fabp4* contain promoter-anchored chromatin looping events that show chromosomal organization changes that correlate with transcriptional changes during adipogenesis(20).

Pparγ2 is a lineage-determining transcription factor that is induced early during adipogenic differentiation(21). Based on the previously reported role for Prmt5 in mediating chromatin looping at the Pparγ2 gene locus(9), we sought to determine whether Prmt5 plays a broader role of Prmt5 in genome-wide higher order chromatin organization and transcriptional control at the onset and into the early stages of adipogenesis using 3T3-L1 cells, a well-characterized cell line for studying adipogenesis. Prmt5 binding sites were abundant in 3T3-L1 cells at the onset of differentiation. Prmt5 localized to thousands of transcriptionally-active regions, where it acted as either a positive or negative modulator of transcription. Prmt5 co-localized with regulators of chromatin organization Ctcf, Smc1 and Med1 at hubs of chromatin looping. Knockdown of *Prmt5* caused significant weakening of TAD boundaries near sites of Prmt5 and Ctcf colocalization. Prmt5-mediated looping events impacted insulation strength, which correlated with the gene dysregulation observed at genes at or in proximity to TAD boundaries upon *Prmt5* knockdown. Altogether, this genome-wide study identifies a broad role for Prmt5 in transcriptional control and chromatin organization during adipogenesis.

## RESULTS

### Abundant Prmt5 binding to chromatin across the genome at early stages of adipogenesis

Previously we reported that Prmt5 is required for promoter-enhancer and promoter-promoter interactions during adipogenesis at the Pparγ2 locus(9,18). In particular, we found that specific genomic sequences involved in chromatin looping were bound by Prmt5(9). To better understand the roles of Prmt5 during adipogenesis, we performed chromatin immunoprecipitation paired with high-throughput sequencing (ChIP-Seq) in 3T3-L1 cells. Previously published findings from our lab indicated that Prmt5 expression is stable during differentiation(8) but its mediation of long-range interactions is dynamic within the first 48 hours of differentiation(9,18). Therefore, we performed Prmt5 ChIP-Seq at time points at the onset of differentiation (day 0) and on days 1 and 2 of 3T3-L1 cell differentiation. Read numbers and correlation of replicates are reported in Supplemental Table 1.

Initial assessment of Prmt5 binding revealed pronounced Prmt5 promoter occupancy (Figure 1A). The *alpha beta hydrolase domain protein 2* (*Abhd2*) promoter contained the strongest Prmt5 peak in the dataset, with a narrow, defined peak resembling a transcription factor peak. Broader Prmt5 peak distribution was observed at other genomic sites such as the promoter for *platelet derived growth factor receptor alpha* (*Pdgfra*), which encodes a preadipocyte marker (Figure 1B). Annotation of day 0 Prmt5 peaks showed a large proportion of peaks located within or in proximity to gene bodies, with 19% of Prmt5 peaks at promoter-TSS sites and ∼45% associated with gene bodies (Figure 1C). Similar annotation patterns were observed for day 1 and day 2, with a modest increase in promoter-TSS localization and modest decreases in the number of intron and intergenic peaks (Supplemental Figure 1A). Overall, 38,259 significant Prmt5 peaks were detected at day 0, while 8,374 peaks were detected at day 1, and 6,285 peaks at day 2 (Supplemental Table 2). Gene Ontology (GO) analyses revealed that at all three time points, Prmt5 was binding to genomic regions annotated to regulators of gene expression and regulators of RNA processing (Figure 1D, Supplemental Figure 1A), consistent with known roles for Prmt5 in gene regulation and in splicing(15). Motif analysis of Prmt5 peaks revealed that the Nrf1 binding motif was uniquely observed to be enriched at Prmt5-bound chromatin sites at all time points (Figure 1E, Supplemental Figure 1A). Nrf1 is a transcription factor associated with genes controlling metabolism and mitochondrial respiration, and its importance in adipose differentiation and function has been reviewed(22). When examining Prmt5 occupancy changes with differentiation, we observed that the peak enrichment score at the *Abhd2* and *Pdgfra* promoters was lower at day 1 and day 2, though still well above background (Figures 1A-B). Genome-wide assessment of Prmt5-binding in the proximity of gene loci showed relatively stable Prmt5 binding between day 0, day 1, and day 2 of differentiation (Figure 1F, Supplemental Figure 1B). K-means clustering identified 3 patterns of Prmt5 binding that correlate with stable gene expression based on analysis of existing RNA-seq data from 3T3-L1 differentiation experiments(20) (Figure 1F). Tag density scores at gene promoters are similar for clusters 1, 2 and 3. Cluster 1 has the narrowest Prmt5 peaks on average while cluster 2 and 3 have broader Prmt5 binding sites.

**Figure 1.**
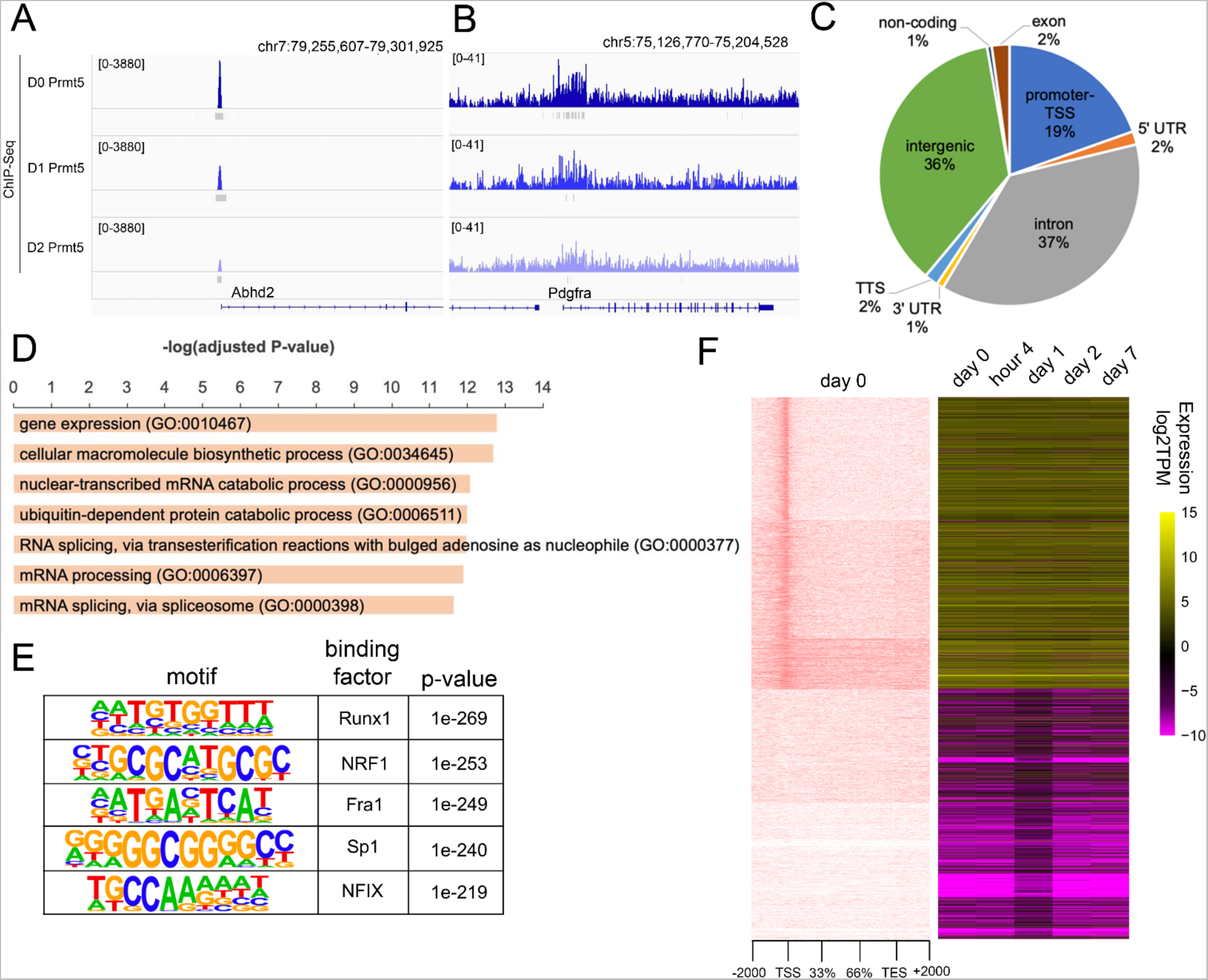
Genome-wide binding patterns for Prmt5. (A) Genome Browser tracks showing Prmt5 ChIP-seq at day 0, day 1, and day 2 of 3T3-L1 differentiation. Significant ChIP peaks determined by MACS2 are indicated below the genome browser tracks in gray for the *Abhd2* promoter and (B) the *Pdgfra* promoter. (C) Gene annotation of all significant Prmt5 peaks at day 0. (D) Top biological processes identified from Gene Ontology analysis of significant day 0 Prmt5 peaks. (E) The HOMER motif discovery algorithm identified Runx1 and Nrf1 as the top motifs present within Prmt5 peaks. (F) Prmt5 ChIP-Seq tag density plots over the gene body ± 2 kb displayed as heat maps, order determined by k-means clustering. Expression data displayed as log_2_TPM (transcripts per million mapped reads) for day 0, day 1, day 2, and day 7 of 3T3-L1 differentiation.

### Prmt5 binds to transcriptionally active regions of the genome

Prmt5 binding in the vicinity of gene bodies was highly correlated with active transcription of the gene (Figure 1F). Our Prmt5 ChIP-Seq data validated previously published findings showing inducible Prmt5 binding at the *Pparγ2* promoter and -10kb enhancer(9) (Supplemental Fig 1C - top). The promoter for the key adipogenic regulator *Cebpa* also showed Prmt5 binding (Supplemental Fig 1C - middle). However, Prmt5 occupancy was limited or absent at adipogenic genes that are expressed later in differentiation, such as *Adipoq* (Supplemental Fig 1C - bottom). This finding is consistent with prior data showing inducible Prmt5 binding at phenotypic adipocyte genes that coincides with the activation of those genes at later times of differentiation(8). These data indicate that Prmt5 binding correlates with gene expression at the onset and at early phases of adipogenesis.

To further support this conclusion, we next examined how Prmt5 binding related to the deposition pattern of select histone modifications(23). Tag density plots of histone modifications H3K4me3, H3K27me3 and H3K27ac centered on Prmt5 day 0 peaks revealed a genome-wide overlap of Prmt5 binding with H3K4me3 and H3K27ac (Figure 2A-B), histone markers of transcriptional activity. This pattern was also evident when looking at individual Prmt5 promoter peaks at the *Abhd2* promoter (Supplemental Figure 2A) and the *Tom1* promoter (Figure 2C) as well as at Prmt5 intronic peaks such as that in the *Adipor2* gene (Supplemental Figure 2B). Quantification of histone and Prmt5 ChIP-Seq overlap showed that 53% (20,396) of the total 38,259 day 0 Prmt5 peaks overlapped with H3K27ac while 37% (14,155) overlapped with H3K4me3 peaks (Figure 2C). At day 2, 70% (4,411) of Prmt5 peaks overlapped with H3K27ac and 59% (3,677) of Prmt5 chromatin binding sites overlapped with H3K4me3 sites. Minimal overlap between Prmt5 and H3K27me3 peaks was observed at day 0 and 2 (Figure 2D). This assessment of the epigenomic landscape indicates that Prmt5 localizes to regions that are transcriptionally active.

**Figure 2.**
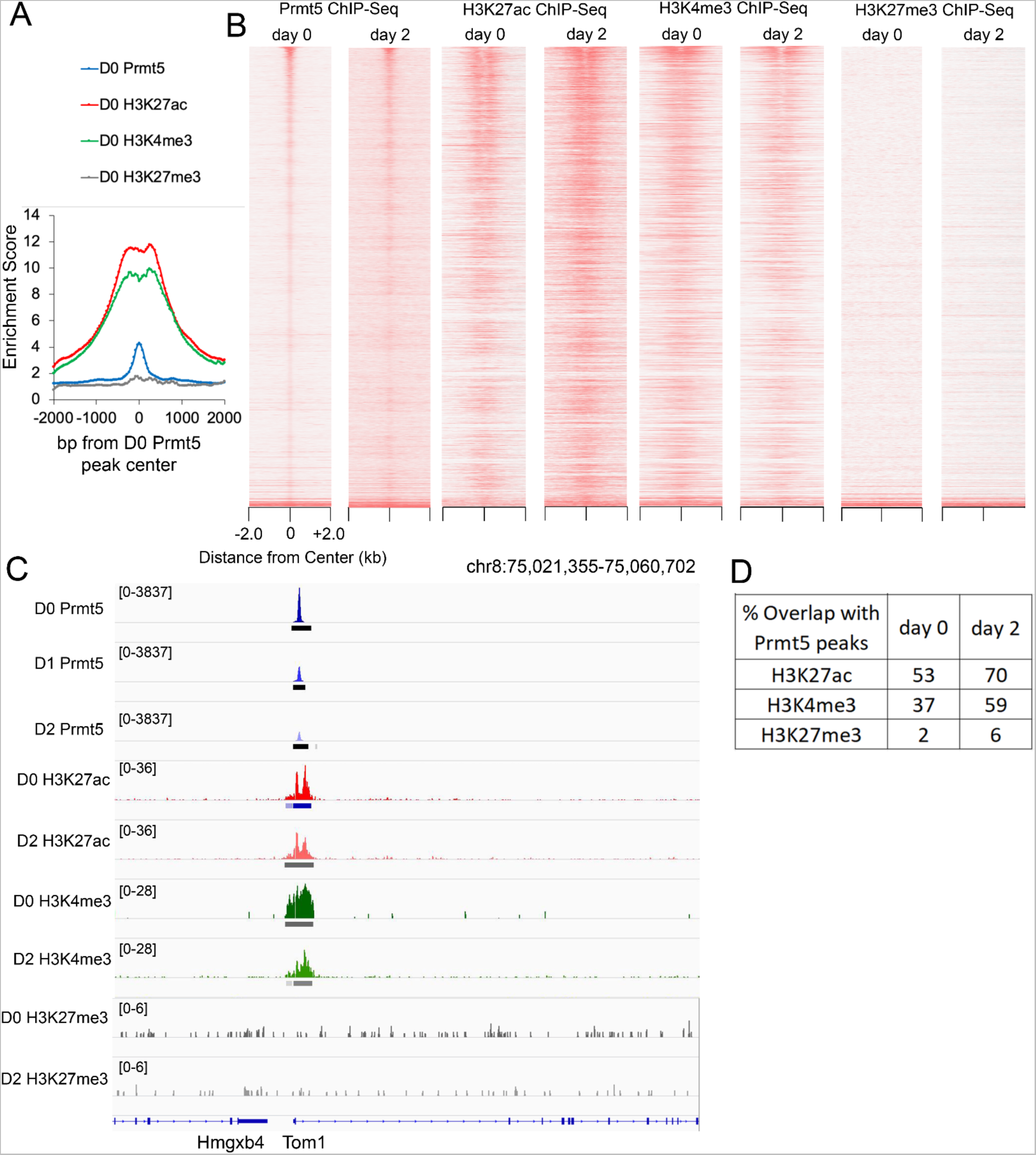
Association between histone modifications and Prmt5 binding. (A) Average tag density plots for Prmt5, H3K27ac, H3K4me3, and H3K27me3 ChIP-Seq, in relation to Prmt5 peak centers. (B) Tag density plots for day 0 and day 2 ChIP-Seq for Prmt5, H3K27ac, H3K4me3, and H3K27me3, plotted +/- 2kb of Prmt5 peak centers. (C) Genome Browser tracks showing Prmt5 ChIP-seq at day 0, day 1, and day 2 of 3T3-L1 differentiation, along with the corresponding tracks for H3K27ac, H3K4me3, H3K27me3 ChIP-Seq at day 0 and day 2 of 3T3-L1 differentiation. Significant ChIP peaks determined by MACS2 are indicated below the genome browser tracks. (D) bedtools intersect was used to calculate significant peak overlap between Prmt5 ChIP-Seq and histone modification ChIP-Seqs at day 0 and day 2 of differentiation.

### *Prmt5* knockdown causes both positive and negative changes in gene expression

To gain insight into the functional significance of the Prmt5 binding patterns, we performed siRNA-mediated knockdown of *Prmt5* using previously validated siRNAs for *Prmt5*(9). When siRNA transfection was performed in proliferating cells, 50-70% knockdown was observed (Supplemental Figure 3A), consistent with prior studies(9). When cells treated with *Prmt5* siRNA were induced to differentiate at day 0, RT-qPCR analysis of *Prmt5* mRNA showed a decrease in *Prmt5* expression at day 0, hour 6, day 1, and day 2 of differentiation (Supplemental Figure 3B). Consistent with prior data(8), *Prmt5* knockdown blocked upregulation of the early adipogenic factors *Pparγ2* and *Cebpa* (Supplemental Figure 3B), genes critical for successful progression through adipogenesis. This downregulation of Prmt5 was also observed by Western blot (Supplemental Figure 3C). Prior studies showed that a decrease in Prmt5 at the onset of adipogenesis in multiple cell culture models prevents differentiation(8,9,18).

We examined genes whose transcription appeared to be dependent on Prmt5 chromatin binding over the initial course of 3T3-L1 differentiation. Three genes, *Ptn*, *Thbs2* and *Pdgfra*, were identified as having Prmt5 promoter binding at day 0 of differentiation but no or decreased binding by day 2 (Figures 1B, 3A). *Ptn* and *Thbs2* are known negative regulators of adipogenesis(24,25) while *Pdgfra* is a known preadipocyte marker(26,27). Their transcription is normally downregulated with induction of differentiation (confirmed in Figure 3B). *Ptn*, *Thbs2*, and *Pdgfra* expression were significantly reduced when Prmt5 was knocked down (Figure 3B), indicating that Prmt5 is a transcriptional co-activator of adipogenic repressors in undifferentiated cells. The data reflect complexity in the role of Prmt5 as a transcriptional regulator at different times of the differentiation process as it coactivates genes associated with the precursor state prior to differentiation and coactivates genes associated with the differentiated state during early adipogenesis.

**Figure 3.**
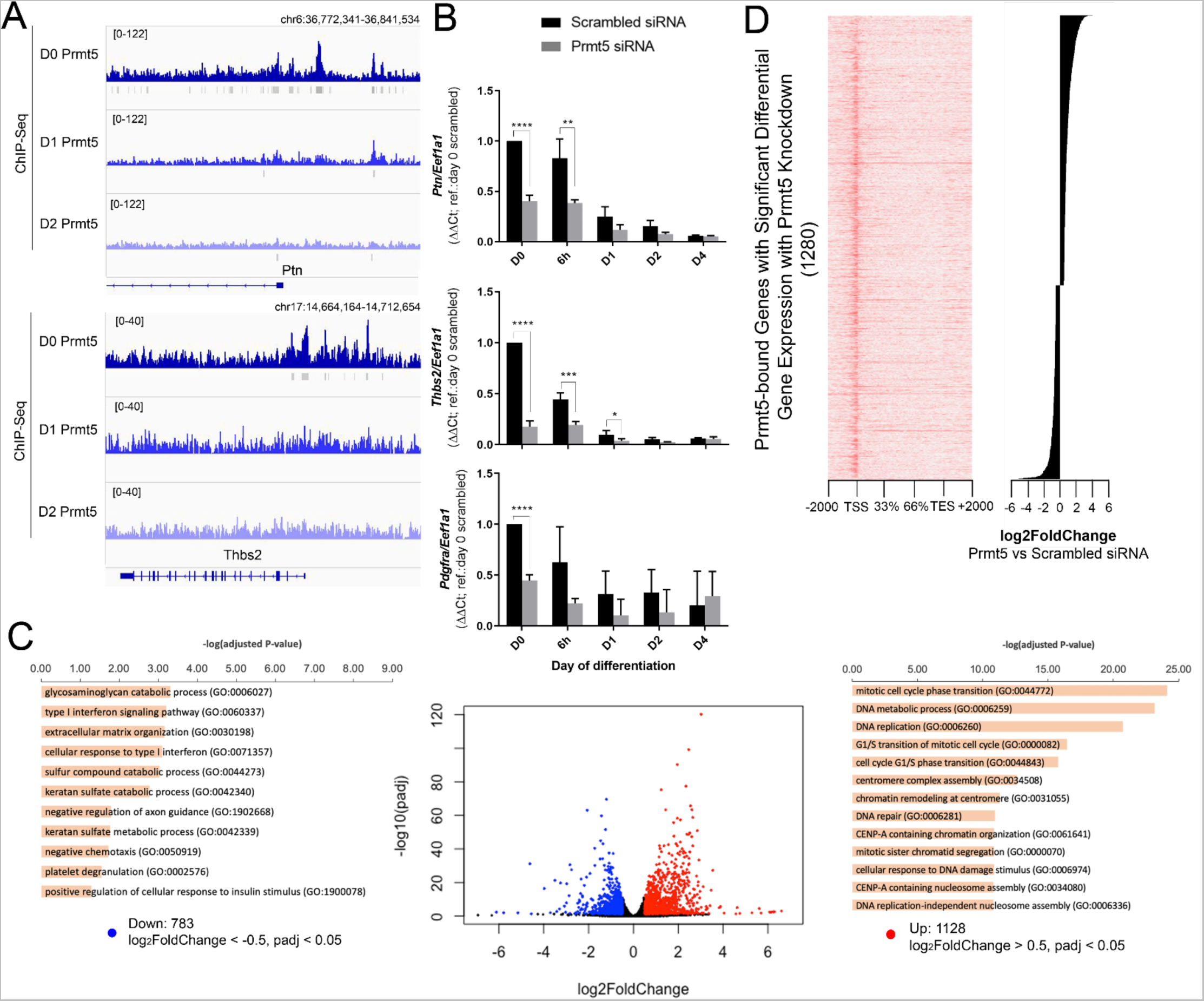
Prmt5-dependent gene expression changes. (A) Genome Browser tracks showing Prmt5 ChIP-seq at day 0, day 1, and day 2 of 3T3-L1 differentiation. Significant ChIP peaks determined by MACS2 are immediately below the genome browser tracks in gray. (B) Real-time RT-qPCR analysis of Prmt5-bound genes in 3T3-L1 cells transfected with scrambled or *Prmt5*-targeting siRNA. Cells were collected at time points between D0 and D4 of differentiation. Data represent averages from 3-6 replicates and are presented as means +/- standard deviations (SD). The level of expression in scrambled day 0 samples were set to a value of 1 for genes *Ptn*, *Thbs2*, and *Pdgfra*, and other values are relative to that sample value. *, P≤0.05; **, P≤0.001; ***, P≤ 0.0001 (versus scrambled by Student’s t test). (C) Volcano plots displaying differentially expressed genes between scrambled and *Prmt5* siRNA transfected 3T3-L1 cells harvested at day 0 of differentiation. The y-axis data correspond to the mean log10 expression levels (padjusted-values). The red dots represent significantly upregulated genes (padj < 0.05, log_2_foldchange > 0.5) and blue dots represent significantly downregulated genes (padj < 0.05, log_2_foldchange < −0.5). The black dots represent genes whose differential expression levels did not reach statistical significance. Flanking the volcano plot are Gene Ontology Biological processes charts for the top upregulated and downregulated genes. (D) Differential expression of Prmt5-bound Prmt5- dependent genes ranked by log_2_foldchange displayed on the right, with corresponding day 0 Prmt5 ChIP-Seq tag density plots over the gene body ± 2 kb displayed to the left.

To better understand this complexity of function, we identified Prmt5-regulated genes transcriptome-wide via RNA-Seq in Day 0 cells transfected with siRNA targeting Prmt5. Prmt5 knockdown resulted in significant upregulation of 1,128 genes (padj < 0.05, log_2_foldchange > 0.5) and significant downregulation of 783 genes (padj < 0.05, log_2_foldchange < −0.5) (Figure 3C, Supplemental Table 3). Gene Ontology analysis of downregulated genes did not reveal any specific biological processes with padj < 0.001. Examples of lower confidence biological processes include glycosaminoglycan catabolic processing and type 1 interferon signaling pathway (Figure 3C). However, we did note an additional adipogenic inhibitory gene, *Dlk1*(28), which encodes the protein Pref1, to have significant transcriptional downregulation with Prmt5 knockdown (Supplemental Table 3). These observations highlight the finding that Prmt5 coactivates adipogenic inhibitors important for maintaining the preadipogenic state. Gene Ontology analysis of genes significantly upregulated by Prmt5 knockdown indicated that Prmt5 may be important for repressing regulators of cell cycle phase transition (Figure 3C). In line with previously published studies, Prmt5 repressed promitotic genes, *CycB* and *Bub1b*(1). The results indicate that while Prmt5 is associated with transcriptionally active chromatin, it can modulate gene expression of actively transcribed genes both positively and negatively. When examining the 1911 genes differentially expressed with Prmt5 knockdown, 1,280 genes were found to have Prmt5 chromatin binding within their gene body (Figure 3D). These findings suggest that Prmt5 chromatin binding may directly regulate the transcription of ∼66% of genes showing differential gene expression with Prmt5 knockdown. Figure 3D displays Prmt5 bound genes showing differential expression upon Prmt5 knockdown relative to the extent of differential expression.

It is important to note that Prmt5-dependent regulation of gene transcription represents a small fraction of all Prmt5 binding sites genome-wide. Specifically, of the 24,420 day 0 Prmt5 peaks localized to the promoter or gene body, only 1,999 (8.2%) of these peaks corresponded to genes that showed significant transcriptional change upon Prmt5 knock down. This strongly suggests that chromatin binding by Prmt5 is affecting other processes in addition to transcriptional control. Since our previously published work suggested a role for Prmt5 in chromatin looping, we next investigated whether Prmt5 chromatin binding was associated with higher-order genome organization and whether this relates to Prmt5-mediated transcriptional control.

### Prmt5 co-localizes with mediators of genome organization at chromatin loop anchors

Chromatin looping is a phenomenon critical for shaping higher-order genome structure(29,30). Chromatin looping events can occur within and across TADs and can be dependent on Ctcf, Cohesin, the Mediator complex, RNAPII, and/or cell-specific transcription factors(19,31–34). Looping events can also occur between promoters and enhancers(34,35). Integration of previously published promoter Capture Hi-C data in day 0 3T3-L1 cells(20) with our ChIP-Seq data identified Prmt5-anchored chromatin looping events at the *Pdgfra*, *Abhd2*, *Cebpa* and *Cebpg* promoters (Figure 4A, Supplemental Figure 4A). These promoters showed co-localization of Prmt5 with H3K27ac and H3K4me3, while the *Pdgfra* and *Cebpa* promoters also showed Prmt5 colocalization with chromatin organizers Med1, Smc1, and Ctcf. We next examined whether Prmt5 binding coincides with genome-wide binding of Med1, Smc1, and Ctcf. Tag density plots centered on Prmt5 day 0 peaks showed a high coincidence of Prmt5 binding with Med1 and Smc1 (Figure 4B-C). Statistically assessment of the peaks with using MACS2 and quantification of peak overlaps showed that 39% of day 0 Prmt5 peaks overlapped with Med1, 66% with Smc1, and 13% with Ctcf (Figure 4D). Day 2 Prmt5 peaks showed similar extent of overlap with the peaks of these chromatin organizing proteins, suggesting that the associations were relatively stable between day 0 and day 2 (Figures 4C-D), despite the introduction of adipogenic signaling that induces differentiation.

**Figure 4.**
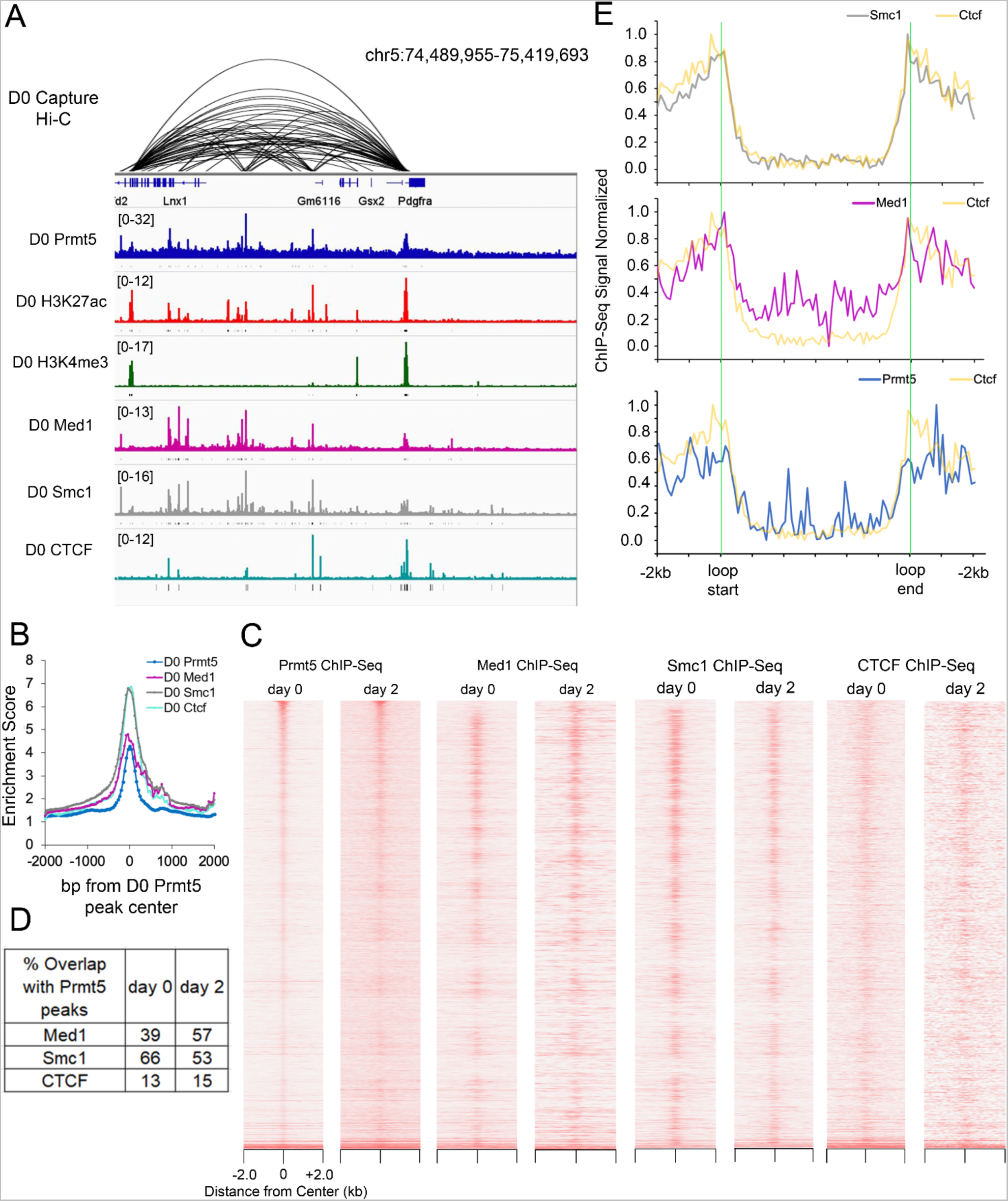
Co-localization of Prmt5 with loop anchors and regulators of genome structure. (A) Genome Browser tracks for showing Prmt5 ChIP-seq at day 0 of 3T3-L1 differentiation, along with the corresponding tracks from H3K27ac, H3K4me3, Med1, Smc1, and Ctcf ChIP- Seqs with D0 Promoter Capture Hi-C displayed above. Significant ChIP peaks determined by MACS2 are indicated below the genome browser tracks. (B) Average tag density plots for Prmt5, Ctcf, Med1, and Smc1 ChIP-Seq, in relation to Prmt5 peak centers. (C) Tag density plots for day 0 and day 2 ChIP-Seq for Prmt5, Med1, Smc1, and Ctcf, plotted +/- 2kb of Prmt5 peak centers. (D) bedtools intersect was used to calculate significant peak overlap between Prmt5 ChIP-Seq and Med1, Smc1, and Ctcf ChIP-Seqs at day 0 and day 2 of differentiation. (E) Average tag density plots showing Smc1, Med1 and Prmt5 binding at DNA loop anchors +/-2kb, in relation to Ctcf binding.

We also asked whether regions of overlap between Prmt5 and modulators of chromatin organization coincide with chromatin loop anchors. We used a Random Forest classification framework for predicting chromatin loops from genome-wide contact maps(36) based on a previously published day 0 3T3-L1 Hi-C dataset(20). This analysis predicted 15,685 high-confidence chromatin loops. As expected, loop boundaries showed a high coincidence of Smc1, Ctcf, and Med1 binding (Figure 4E). Furthermore, the Ctcf binding motif was the most significant motif present (Supplemental Figure 4B). We then examined Prmt5 binding at these loop anchors (Figure 4E). We found a high frequency of Prmt5 binding at loop boundaries; when quantified, 4,989 of 15,685 looping events (31.8%) had Prmt5 binding at one or both loop boundaries.

### *Prmt5* knockdown impacts TAD boundary strength and gene expression

The association between Prmt5 binding and chromatin organizers raised the question of whether Prmt5 plays a regulatory role in the formation of chromatin structures. To address this question, we performed Hi-C in day 0 3T3-L1 cells transfected with scrambled or Prmt5 siRNA and integrated the results with our ChIP-Seq and RNA-Seq data. Read numbers and correlation of replicates are reported in Supplemental Table 4. Eigenvector decomposition analysis to determine the positions of A and B compartments showed no change upon *Prmt5* knockdown (Figure 5A). We performed insulation score analysis to identify TAD boundaries. Insulation scores reflect the relative frequency of interactions across specific genomic bins for a preset genomic span around that bin. Insulation scores were assigned to 10 kb genomic intervals along each chromosome; minima were defined as TAD boundaries, as previously described(37). Of the 4,128 predicted TAD boundaries for scrambled and *Prmt5* knockdown, 87.6% were invariant. Insulation analysis plots show some invariant TAD boundaries at specific genomic regions (Figure 5B). Prmt5-bound chromatin looping events showed considerable insulation strength differences between scrambled and *Prmt5* knockdown (Figure 5B). Prmt5-bound genes *Fzd1* and *Cdk14* are flanking and located within a TAD boundary, respectively. Both genes were downregulated upon *Prmt5* knockdown (*Fzd*1 log_2_foldchange= −1.24, padj= 3.79E-09; *Cdk14* log_2_foldchange= −0.44, padj= 8.13E- 07). This down-regulation appears to correspond with strengthening of the TAD boundary and increased interaction frequency within the adjacent TADs (Figure 5B – top). Conversely, *Prmt5* knockdown caused significant *Myc* gene upregulation (log_2_foldchange= 0.73, padj= 9.56E-12), which corresponded with a weakening of the TAD boundary encompassing the gene (Figure 5B – bottom). These findings reveal that Prmt5 knockdown can impact TAD boundary strength and suggest that Prmt5-mediated chromatin looping events play a role in modulating TAD boundary strength, which in turn impacts gene regulation.

**Figure 5.**
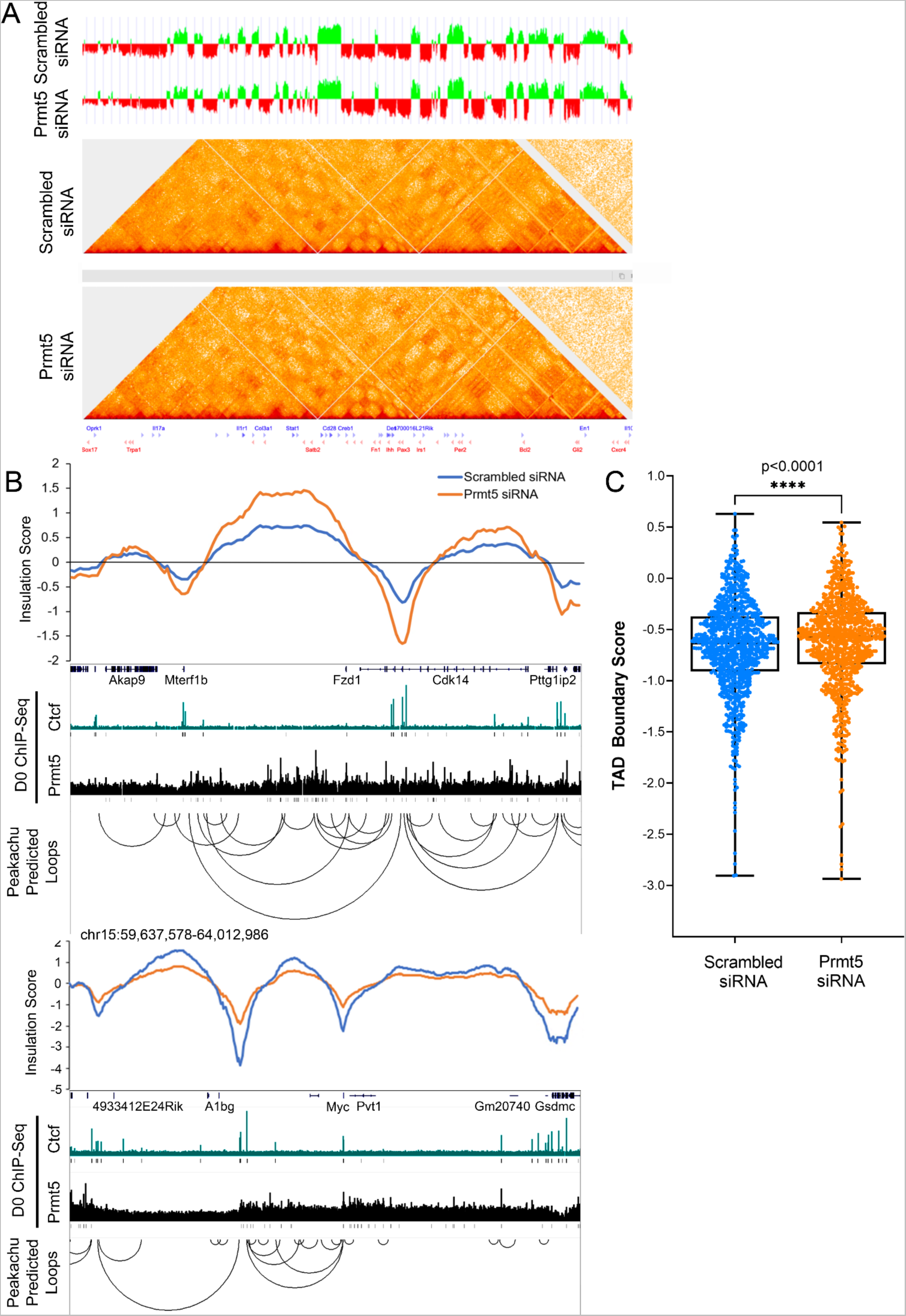
Effect of *Prmt5* knockdown on TADs and TAD boundaries. (A) Compartment profiles (the first principal components) of scrambled siRNA and *Prmt5* siRNA data for Chr 1. The A-type (open) compartments are shown in green, and the B-type (closed) compartments are shown in red. Below this, contact heatmaps are displayed. (B) Insulation plot profiles at 10-kb intervals are displayed genomic loci around genes *Cdk14* and *Myc* for scrambled siRNA and *Prmt5* siRNA transfected samples. Below that, a genome browser track for the same locus is displayed showing Prmt5 and Ctcf ChIP-seq peaks at day 0 of 3T3-L1 differentiation. High-confidence D0 3T3-L1 looping event interactions are depicted as purple lines below this plot. (C) Insulation scores were plotted for all Prmt5-bound TAD boundaries for day 0 scrambled siRNA and *Prmt5* siRNA transfected samples.

To more broadly determine whether there is a connection between Prmt5-dependent gene expression and TAD boundary strength changes, we looked at those 1,280 Prmt5- bound genes with significant differential gene expression described in Figure 3D and examined their proximity to a TAD boundary. 20.4% of these differentially expressed genes were found spanning a predicted TAD boundary or were within 50kb of one. We examined insulation strength at all Prmt5-bound TAD boundaries (Figure 5C). This analysis shows overwhelming evidence that *Prmt5* knockdown can contribute to significant TAD boundary weakening (a more positive insulation score). This provides a possible mechanism for the transcriptional dysregulation observed with Prmt5 knockdown.

CTCF-anchored chromatin loops are very frequently found at TAD boundaries(31). To support our finding that genes affected by *Prmt5* knockdown are predominantly found at TAD boundaries, we examined how many *Prmt5*-knockdown differentially expressed genes (Prmt5-KD DEG) contain Prmt5-bound chromatin sites that colocalize with CTCF. Genome wide analysis revealed that 402 of our Prmt5-KD DEGs (32%) contain Prmt5 ChIP peaks that colocalize with CTCF. This colocalization rate is considerably higher than the genome-wide incidence of Prmt5 colocalization with CTCF (13%, Figure 3D). Our findings suggest that Prmt5 knockdown may impact TAD boundary strength at Prmt5-dependent loci located at or near CTCF-bound TAD boundaries leading to gene dysregulation.

## DISCUSSION

Prmt5 is a versatile protein that regulates numerous cellular processes. In the cytoplasm, Prmt5-mediated methylation of Sm proteins impacts spliceosome assembly and RNA splicing(38–41) while Prmt5-mediated methylation of p53 in the nucleus is important for p53-mediated G1 arrest(42,43). Prmt5 has also been implicated in direct transcriptional control, with Prmt5-mediated histone methylation determining transcriptional activation and repression in a context-dependent manner. However, genome-wide ChIP-Seq studies examining Prmt5-mediated enrichment of H4R3me2s revealed no correlation with gene expression(44), strongly suggesting that Prmt5 also may have a role in the nucleus that is distinct from transcriptional control. That study also implicated Prmt5 in chromosomal organization due to observations that H4R3me2s colocalizes to chromatin regions bound by Smc1 and the Mediator complex(44). This, in conjunction with our own studies linking Prmt5 chromatin binding to promoter-enhancer looping at the *Pparγ2* locus and looping events between adipogenic loci(9,18), suggested a role for Prmt5 in chromatin organization.

In the present study, we took a genome-wide approach to explore patterns of Prmt5 binding from the onset of adipogenesis through day 2, when adipogenic lineage determinants that initiate the adipogenic program are induced. Based on the large number of Prmt5 peaks at day 0, we also examined Prmt5-mediated transcriptional control and chromatin organization at the onset of differentiation. Prmt5 ChIP-Seq allowed us to examine chromatin binding dynamics during adipogenesis. Between day 0 and day 1 of differentiation, more than 30,000 Prmt5 peaks were lost, suggesting an important role for Prmt5 chromatin binding in the undifferentiated preadipocyte state and dynamic regulation of Prmt5 binding as adipogenesis progresses. Prmt5 knockdown studies in day 0 3T3-L1 cells identified numerous adipogenic repressors that are transcriptionally activated by Prmt5, including *Ptn*, *Thbs2*, *Dlk1*, and *Pdgfra*. Our new findings strongly suggest that Prmt5 is critical for regulating preadipocyte identity genes that are critical for maintaining the preadipocyte state and that must undergo downregulation to proceed through adipogenesis.

While previously we had reported that Prmt5 knockdown can block adipogenesis, we had attributed this primarily to Prmt5-dependent up-regulation of the master regulator of adipogenesis, Pparγ2. ChIP-Seq and RNA-Seq integration show that Prmt5 is a critical inhibitor of a number of cell cycle regulators, including CycB and Bub1b(1). Our studies point to the targeting of cell cycle regulators as an additional reason why Prmt5 knockdown blocks adipogenesis. Prmt5 is a critical factor during other developmental processes, including embryonic development, erythrocyte differentiation, myogenesis, hematopoiesis, and neural stem cell differentiation(1–4). Follow-up investigations examining whether Prmt5 regulates cell cycle regulators in other adult stem cell populations may further elucidate why Prmt5 plays such a critical role in adipogenesis and development.

While we provide evidence of Prmt5-mediated transcriptional control, this represents a relatively small proportion of the 38,259 Prmt5 chromatin binding sites identified in day 0 3T3-L1 cells. In conjunction with H4R3me2s genome-wide binding studies that showed no correlation to transcription, we can conclude that Prmt5 is not solely a transcriptional regulator in the nucleus. Our study revealed colocalization of Prmt5 with Smc1 and Med1 and with a smaller subset of peaks co-localizing with CTCF. We also made the novel observation that Prmt5 can bind to chromatin looping anchors across the genome. These findings strongly suggest that Prmt5 has a previously unidentified role in chromatin organization. Our Hi-C experiments with Prmt5 knockdown resulted in TAD boundary strength changes at select loci where Prmt5 colocalizes with CTCF. Prmt5 may be a critical chromatin loop anchor protein for chromosomal organization and Prmt5 may also be important for establishing regions of high insulation that characterize TAD boundaries. Histone modifiers have been implicated in genome organization before,(45–47) and Prmt5 may be another novel example of how histone modifiers are important for shaping the genome architecture.

Chromatin organization undergoes dynamic changes during stem cell differentiation and there is a great deal of interest in understanding how these changes impact transcription. Genome reorganization has been suggested as a mechanism for transcriptional change in differentiation processes as well as disease(48,49). While *Prmt5* knockdown only affected the expression of a small subset of Prmt5-bound genomic regions, our data suggest that Prmt5-mediated chromatin looping events may be contributing to TAD boundary strength and this may be a potential mechanism by which Prmt5 regulates gene transcription. Further work will be required to understand the mechanistic details connecting the effects of Prmt5 on genome architecture and its impact on transcription.

Our data showed that Prmt5 can both activate and repress transcription, depending on the context. The fact that this regulation is accomplished in transcriptionally active regions of the genome devoid of H3K27me3 and H3K9me3 suggests that Prmt5 is fine-tuning active gene expression. In support of this idea, recent studies modeling the impact of post-translational modifications (PTM) on chromatin contact frequency have indicated that PTM deposition changes at select promoters can lead to transitions between high- and low- transcribing states(50). Prmt5-mediated enhancer-promoter contacts may be a mechanism that selects genes for repression or activation. These contacts may also be important for establishing regions of high insulation throughout the genome. A significant finding was the enrichment of genes that are bound and regulated by Prmt5 at or near TAD boundaries. This suggests that Prmt5 functions as a transcriptional regulator only in specifically structured regions of the genome and raises the possibility that chromatin-bound Prmt5 in other regions of the genome has other roles that remain to be defined.

Many interesting questions remain. It will be important to investigate how looping affects gene expression and whether Prmt5-dependent histone arginine methylation is a contributing factor. It will also be important to examine how Prmt5 is recruited to its chromatin targets and how its recruitment affects the surrounding epigenomic landscape. Prior studies have shown that Prmt5-mediated histone methylation is required for recruitment of Wdr5 and the MLL complex to gene targets(51). Further experiments examining chromatin landscape changes in *Prmt5* knockdown cells may reveal additional mechanisms by which Prmt5 is modulating transcription. In summary, our genome-wide studies have identified unique functions for Prmt5 chromatin binding in the nucleus for transcriptional control and for nuclear organization.

## MATERIALS AND METHODS

### Antibodies

For chromatin immunoprecipitation (ChIP) experiments, rabbit polyclonal Prmt5 antibody (Epigentek, catalog # A-3005-100) and normal Rabbit IgG (Cell Signaling Technology, catalog # 2729) were used. For Western Blot, mouse monoclonal Prmt5 antibody (Santa Cruz, catalog # sc-376937) and mouse monoclonal Vinculin antibody (Santa Cruz, catalog # sc-25336) were used.

### Biological Resources

3T3-L1 cells were purchased from ATCC (Manassas, VA).

### Cell Culture

3T3-L1 cells were maintained at sub-confluent densities in Dulbecco’s modified Eagle’s medium (DMEM) supplemented with 10% fetal bovine serum (FBS) and 1% penicillin-streptomycin in a humidified incubator at 37°C and 5% CO2. For induction of differentiation, 2-day post-confluent cells were treated with an adipogenic cocktail (1 μg/ml insulin, 0.25 μg/ml dexamethasone, 0.5 mM IBMX, with 10% FCS). After 48 h, cells were maintained in medium containing 1 μg/ml insulin and 1% penicillin-streptomycin. Cells were collected at 2- days post-confluence (day 0 of induction of adipogenesis), hour 6, day 1 and day 2 of adipogenesis.

### siRNA Transfection

Cells were transfected at 60–70% confluence using the Lipofectamine 2000 (Invitrogen) reagent with 24 nM siRNA (scrambled or Prmt5 siRNA oligo3 or oligo5, see sequence in Supplemental Table 5). Cells were incubated with siRNA in OptiMEM or DMEM (Invitrogen) supplemented with 10% fetal bovine serum (FBS) and 1% penicillin-streptomycin for 6 hours, after which media was replaced with DMEM supplemented with 10% fetal bovine serum (FBS) and 1% penicillin-streptomycin.

### Chromatin immunoprecipitation (ChIP)

ChIP was performed as previously described(52,53) with some modifications. Briefly, cells were cross-linked with 1% formaldehyde (Ted Pella Inc., Redding, CA) for 10 min at room temperature. After quenching of the formaldehyde with 125 mM glycine for 5 min, fixed cells were washed twice with ice-cold phosphate-buffered saline (PBS) supplemented with protease cocktail inhibitor (PCI) (Thermo Scientific, Pierce Protease Inhibitor Tablets, EDTA-free, catalog number A32965) resuspended in cell lysis buffer (10 mM Tris HCl [pH 7.5], 10 mM NaCl, 0.5% NP-40, and protease inhibitors), and incubated on ice for 10 minutes. The nuclei were pelleted, washed with MNase digestion buffer (20 mM Tris HCl [pH 7.5], 15 mM NaCl, 60 mM KCl, 1 mM CaCl_2_ and PCI), and incubated for 20 min at 37°C in the presence of 1,000 gel units of micrococcal nuclease (catalog number. M0247S; NEB, Ipswich, MA) in a 500 ul volume of buffer. The reaction was stopped by adding 500 ul volume of sonication buffer (100 mM Tris HCl [pH 8.1], 20 mM EDTA, 200 mM NaCl, 0.2% sodium deoxycholate, 2% Triton X-100, and PCI). Samples were sonicated for 10-12 minutes (high intensity, 30s on/30s off) in a Bioruptor Pico system (Diagenode, Denville, NJ) at 4°C, and centrifuged at 21,000xg for 5 min. The length of the fragmented chromatin was between 200 and 500 bp as analyzed on agarose gels. Chromatin concentrations were measured using a Qubit 3 fluorometer (Invitrogen). 5 μg chromatin was subjected to immunoprecipitation with the 2 ug Prmt5 antibody or 2 ug IgG antibody as a negative control at 4°C overnight, and immunocomplexes were recovered by incubation with protein A-agarose magnetic beads (Invitrogen) for 3-4 hours at 4°C. Sequential washes of 5 minutes each were performed with ChIP buffer (50 mM Tris [pH 8.1], 10 mM EDTA, 100 mM NaCl, 1% Triton X-100, and 0.1% sodium deoxycholate), ChIP buffer with 0.5 M NaCl, Tris/LiCl buffer (10 mM Tris [pH 8.1], 1 mM EDTA, 0.25 M LiCl_2_, 0.5% NP-40, and 0.5% sodium deoxycholate) and Tris/EDTA buffer (50 mM Tris [pH 8.0] and 10 mM EDTA). Immune complexes were eluted in 100 ul of elution buffer (10 mM Tris HCl [pH 8.0], 10 mM EDTA, 150 mM NaCl, 1% SDS, 5 mM DTT), incubated for 30 minutes at 65°C. Eluates were reverse cross-linked overnight at 65 °C. Following this, samples were treated with 1ul RNase A (0.5 mg/ml, Thermo Fisher) for 1 hour at 37C and 1 ul of Proteinase K (1 mg/ml) for 2 hours at 37°C. DNA was purified using a ChIP DNA Clean & Concentrator kit (Zymo Research, Irvine, CA).

### ChIP-Seq Library Preparation and Analysis

Sequencing libraries were prepared using the ThruPLEX DNA-seq kits (Rubicon Genomics). Libraries were prepared with Prmt5 ChIP or Input DNA and individually barcoded. qPCR library amplification was performed for five to nine cycles. Libraries were sequenced using an Illumina NextSeq High Output instrument with paired-end 75-bp reads. Fastq files were aligned to the mm10 reference genome with bowtie2(54). Reads with a mapq score of 20 or greater were retained using Samtools. PCR duplicates were filtered out using picard. Mapped read numbers are reported in Supplemental Table 1. Reproducibility of biological replicates (n=2) was verified by Pearson correlation analysis using deeptools(55) (Supplemental Table 1). K-means clustering of ChIP-Seq tag density plots were generated using ngs.plot(56). Peak calling was performed using the macs2(57) analysis algorithm. Peak annotation in relation to RefSeq transcription start sites and motif analysis was done using the HOMER suite(58). Gene Ontology of annotated peaks was performed using the gene list enrichment analysis tool EnrichR(59–61). Bedtools intersect(62) was used to assess peak overlap between different ChIP-Seq datasets.

### RT-qPCR Gene Expression Analysis

RNA was extracted using TRIzol reagent (Invitrogen) and the yield determined by measuring optical density at 260 nm (OD260). 1 μg total RNA was subjected to reverse transcription with a QuantiTect reverse transcription kit (Qiagen, Germantown, MD). The resulting cDNA and Fast SYBR green master mix (Applied Biosystems, Foster City, CA) were used for quantitative PCR. Amplification reactions were prepared with 25 ng of cDNA template, 0.3 μM concentrations of primers, and 2X SYBR green master mix. qCPR was performed on a QuantStudio3 (Applied Biosystems) instrument. Primer sequences are included in Supplemental Table 5.

### Hi-C Chromosome Conformation Capture

Hi-C was performed in duplicate using a previously described method(63,64) and using the Arima HiC+ kit, according to the manufacturers protocols (Arima Genomics, Catalog no. A510008). Briefly, flash frozen cross-linked cells were lysed and then digested with DpnII- containing enzyme mix at 37°C. The DNA overhanging ends were then marked with biotin followed by proximal end ligation. DNA was reversed crosslinked and ligation products were purified and then fragmented by sonication to an average size of 200-300 bp and size-selected to fragments of < 350 bp. We then performed end repair, Biotin-enrichment, and dA-tailing followed by Illumina TruSeq adapter ligation. Samples were amplified and the PCR primers were removed. Hi-C libraries were then sequenced PE50 on an Illumina HiSeq 4000.

### Hi-C Data Processing

Distiller-nf, a modular Hi-C mapping pipeline, was used as previously described(65) for mapping FASTQ sequencing files to the mm10 mouse reference genome (available at https://github.com/mirnylab/distiller-nf). Hi-C contact matrices were generated and stored as .cool files(66) (available at https://github.com/mirnylab/cooler) and visualized using HiGlass(67) (available at https://higlass.io/). Resultant ‘.cool’ files were used in downstream analyses using cooltools (https://github.com/mirnylab/cooltools). For downstream analyses using cworld (https://github.com/dekkerlab/cworld-dekker), .cool files were converted to .matrix using cooltools dump_cworld. As previously described(37), insulation boundary, compartment, and scaled distance plot analyses were performed using cworld packages matrix2insulation.pl, matrix2EigenVectors.py and matrix2scaling.pl, respectively. cworld correlateMatrices.pl was used for correlation analysis between biological replicates which showed high reproducibility (Pearson’s correlation coefficient >0.97 for all chromosomes). Peakachu(68), a Random Forest classification framework that predicts chromatin loops from genome-wide contact maps, was used for generating a 3T3-L1 chromatin loop list. To quantify average loop strength changes, Coolpup.py(69) was used to aggregate looping events and enrichment scores were compared between different conditions.

### RNA-Seq Library Preparation and Sequencing

For RNA sequencing, RNA samples were prepared as described above. RNA libraries were prepared by the Beijing Genomics Institute, described previously(70). Briefly, poly-A containing mRNA molecules were purified using oligo-dT beads. RNA was then fragmented and made into cDNA using reverse transcriptase and random hexamer primers. Second strand cDNA synthesis was then performed using DNA Polymerase I and RNase H. After adapter ligation, these double-stranded cDNA libraries were processed for paired-end 100 bp sequencing on the BGISEQ500 platform. High-quality reads were aligned to the mouse reference genome (GRCm38) using the Bowtie2 software (v2.2.5), and gene expression levels were calculated using RSEM (v1.2.12), a software package for estimating gene and isoform expression levels from RNA-Seq data(71). Differential gene expression analysis was performed using R package DESeq2(72). The significantly differentially expressed genes are listed in Supplemental Table 3. Gene ontology term identification was performed using EnrichR. R packages ggplot2 and pheatmap were used for data visualization.

### Code Availability

Code for Hi-C analyses are available at the following links: distiller-nf (https://github.com/mirnylab/distiller-nf), pairtools (https://github.com/mirnylab/ pairtools), cooltools (https://github.com/mirnylab/cooltools) and cworld (https://github.com/dekkerlab/cworld-dekker).

## DATA AVAILABILITY

3T3-L1 Prmt5 ChIP-seq data were deposited at the Gene Expression Omnibus (GEO) database under accession number GSE148117. Day 0 3T3-L1 HI-C data (scrambled and Prmt5 knockdown) were deposited under accession number GSE193009 and RNA-Seq for Day 0 scrambled and Prmt5 knockdown were deposited under accession number GSE193010. A superseries of all datasets was created under accession number GSE208298.

For Figure 1, RNA-seq data from GEO: GSE95533 was used. For Figure 1, ChIP-Seq data from GEO: GSE21365 was used. For Figure 4, ChIP-Seq data from GEO: GSE95533 and GEO: GSE21365 used. In Figures 4 and 5, loop prediction data from day 0 3T3-L1 Hi-C was used from GEO: GSE95533. Figure 4 promoter Capture Hi-C data is from GEO: GSE21365.

