## Supplementary material for "Protein arginine methyltransferase 5 (Prmt5) localizes to chromatin loop anchors and modulates expression of genes at TAD boundaries during early adipogenesis": Supp. Figures 1-4

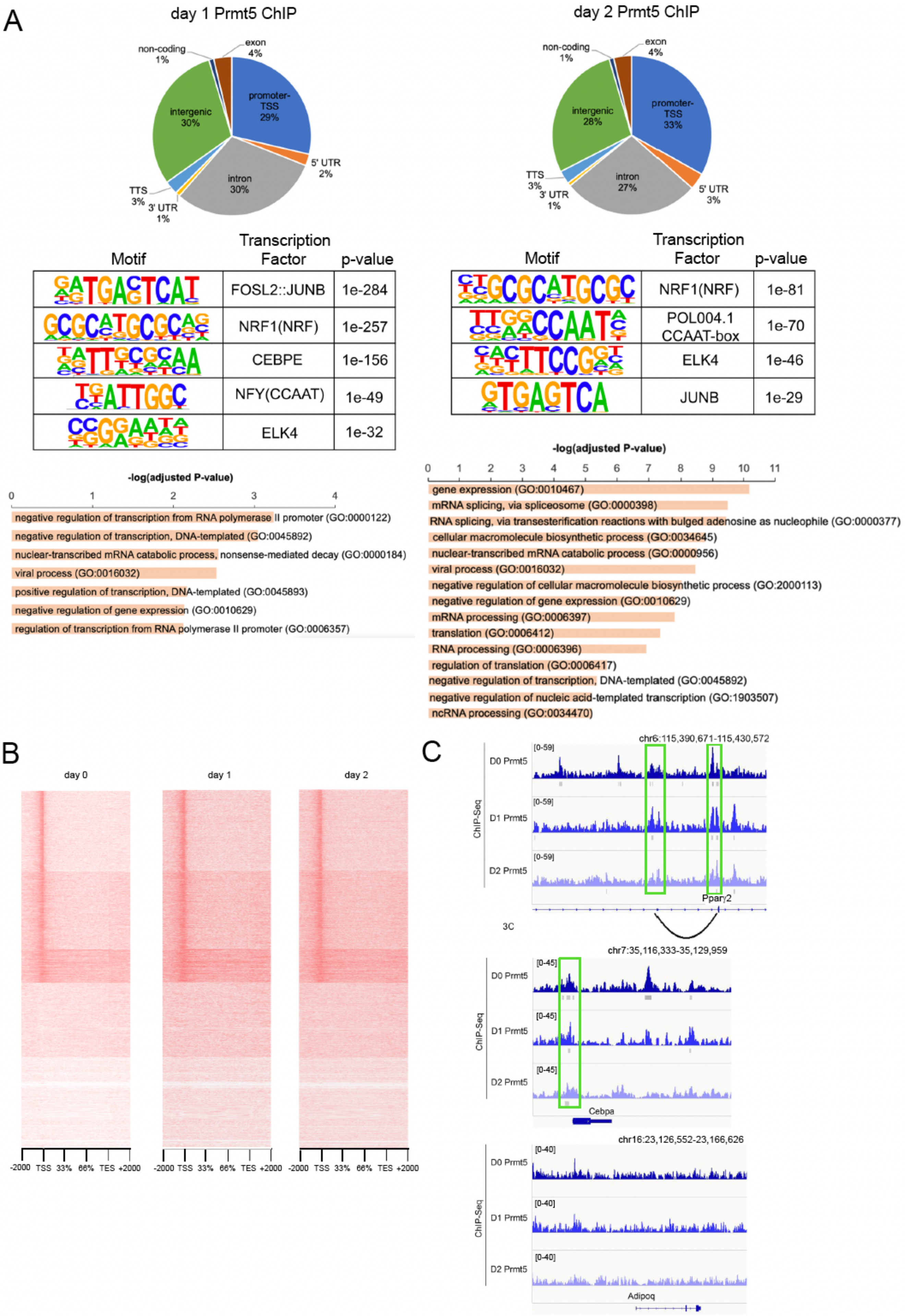
A
day 1 Prmt5 ChIP
non-coding 1%
exon 4%
promoter-TSS 29%
intergenic 30%
5' UTR 2%
intron 30%
TTS 3%
3' UTR 1%
day 2 Prmt5 ChIP
non-coding 1%
exon 4%
promoter-TSS 33%
intergenic 28%
TTS 3%
3' UTR 1%
intron 27%
5' UTR 3%
Motif | Transcription Factor | p-value
FOSL2::JUNB | 1e-284
NRF1(NRF) | 1e-257
CEBPE | 1e-156
NFY(CCAAT) | 1e-49
ELK4 | 1e-32
Motif | Transcription Factor | p-value
NRF1(NRF) | 1e-81
POL004.1 CCAAT-box | 1e-70
ELK4 | 1e-46
JUNB | 1e-29
-log(adjusted P-value)
negative regulation of transcription from RNA polymerase II promoter (GO:0000122)
negative regulation of transcription, DNA-templated (GO:0045892)
nuclear-transcribed mRNA catabolic process, nonsense-mediated decay (GO:0000184)
viral process (GO:0016032)
positive regulation of transcription, DNA-templated (GO:0045893)
negative regulation of gene expression (GO:0010629)
regulation of transcription from RNA polymerase II promoter (GO:0006357)
-log(adjusted P-value)
gene expression (GO:0010467)
mRNA splicing, via spliceosome (GO:0000398)
RNA splicing, via transesterification reactions with bulged adenosine as nucleophile (GO:0000377)
cellular macromolecule biosynthetic process (GO:0034645)
nuclear-transcribed mRNA catabolic process (GO:0000956)
viral process (GO:0016032)
negative regulation of cellular macromolecule biosynthetic process (GO:2000113)
negative regulation of gene expression (GO:0010629)
mRNA processing (GO:0006397)
translation (GO:0006412)
RNA processing (GO:0006396)
regulation of translation (GO:0006417)
negative regulation of transcription, DNA-templated (GO:0045892)
negative regulation of nucleic acid-templated transcription (GO:1903507)
ncRNA processing (GO:0034470)
B
day 0
day 1
day 2
-2000 TSS 33% 66% TES +2000
C
chr6:115,390,671-115,430,572
ChIP-Seq
D0 Prmt5
D1 Prmt5
D2 Prmt5
[0-59]
Pparγ2
3C
chr7:35,116,333-35,129,959
[0-45]
Cebpa
chr16:23,126,552-23,166,626
[0-40]
Adipoq

Supplemental Figure 1. Genome-wide binding patterns for Prmt5. (A) Left panel - Gene annotation, HOMER motif analysis, and Gene Ontology Biological Processes for day 1 Prmt5 ChIP. Right panel - Gene annotation, HOMER motif analysis, and Gene Ontology Biological Processes for day 2 Prmt5 ChIP. (B) Prmt5 ChIP-Seq tag density plots over the gene body ± 2 kb displayed as heat maps for day 0, day 1, and day 2 Prmt5 ChIP-Seq. Order determined by day 0 k-means clustering. (C) Genome Browser tracks showing Prmt5 ChIP-seq at day 0, day 1, and day 2 of 3T3-L1 differentiation for the *Pparγ2*, *Cebpa*, and *Adipoq* genes. The  track indicates the previously reported promoter-enhancer looping event determined by 3C. Significant ChIP peaks determined by MACS2 are immediately below the genome browser tracks in gray.

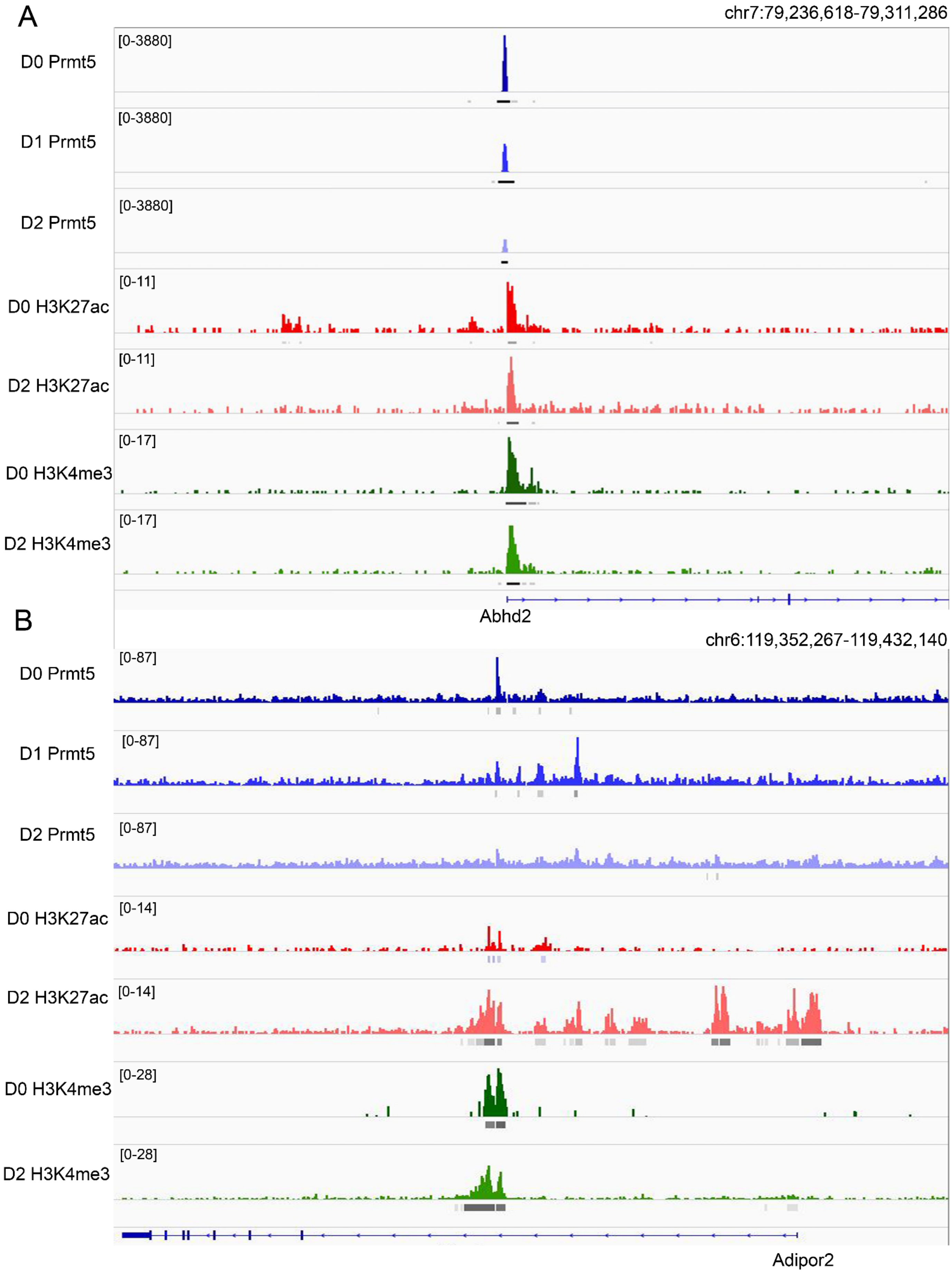
A
chr7:79,236,618-79,311,286
D0 Prmt5
[0-3880]
D1 Prmt5
[0-3880]
D2 Prmt5
[0-3880]
D0 H3K27ac
[0-11]
D2 H3K27ac
[0-11]
D0 H3K4me3
[0-17]
D2 H3K4me3
[0-17]
Abhd2
B
chr6:119,352,267-119,432,140
D0 Prmt5
[0-87]
D1 Prmt5
[0-87]
D2 Prmt5
[0-87]
D0 H3K27ac
[0-14]
D2 H3K27ac
[0-14]
D0 H3K4me3
[0-28]
D2 H3K4me3
[0-28]
Adipor2

Supplemental Figure 2. Association between histone modifications and Prmt5 binding at specific loci. (A) Genome Browser tracks showing Prmt5 ChIP-seq at day 0, day 1, and day 2 of 3T3-L1 differentiation, along with corresponding tracks from H3K27ac, H3K4me3, H3K27me3 ChIP-Seq at day 0 and day 2 of 3T3-L1 differentiation for *Abhd2* and (B) *Adipor2*. Significant ChIP peaks determined by MACS2 are immediately below the genome browser tracks.

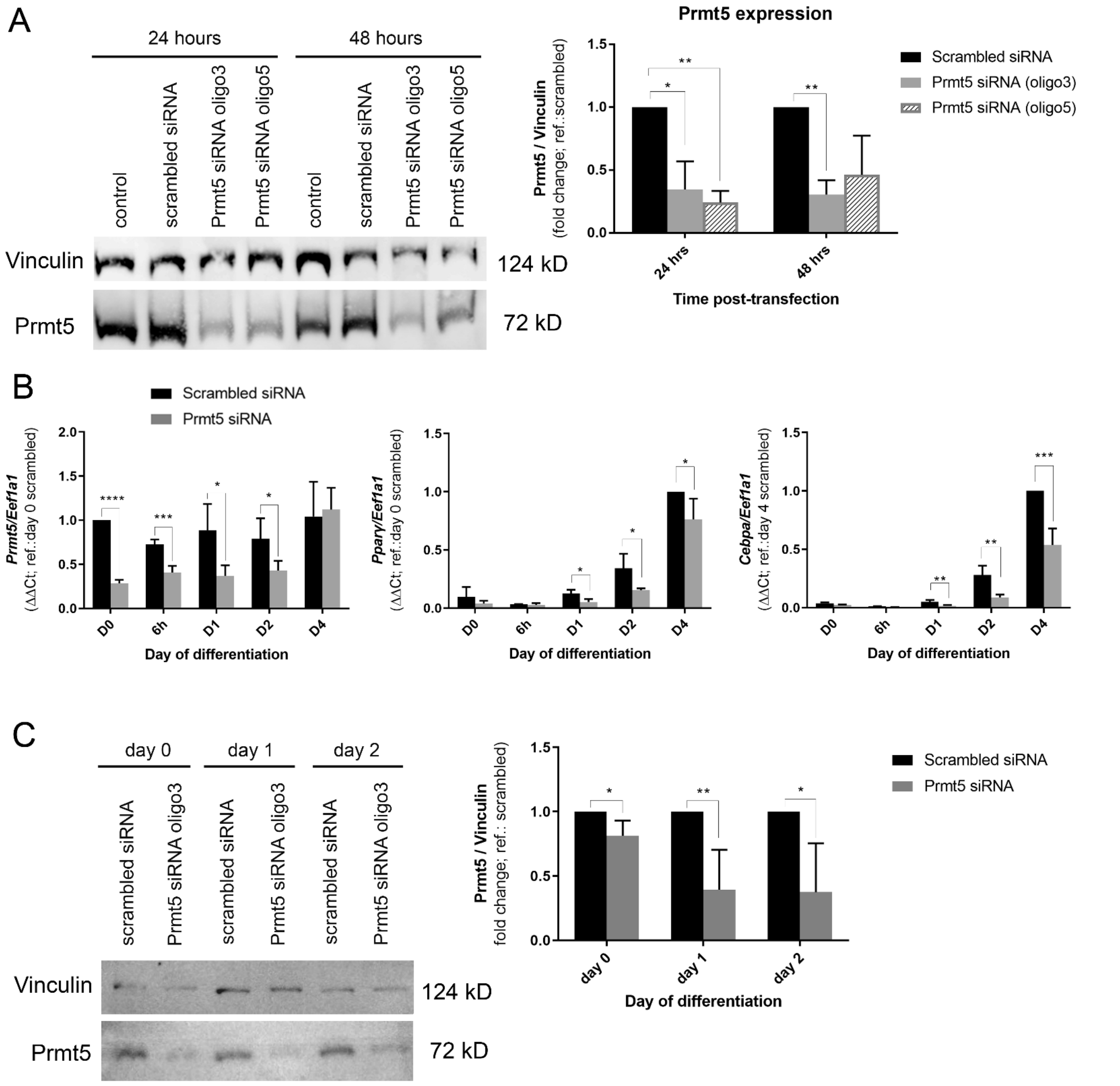


Supplemental Figure 3. *Prmt5* knockdown affects gene expression. (A) Reduced Prmt5 expression detected by Western blotting 24 hours and 48 hours after 3T3-L1 transfection with *Prmt5* siRNAs (oligo3 and oligo5) (means±sd.; n=3/group). P-value calculated in relation to scrambled siRNA samples (*, P≤0.05; **, P≤0.001; ***, P≤ 0.0001) by Student's t-test. (B) Real-time RT-qPCR analysis of *Prmt5*, *Pparγ2*, and *Cebpa*. For genes with inducible expression, *Pparγ2* and *Cebpa*, scrambled day 4 samples were set to a value of 1 and other values are relative to that sample value. Cells were collected at time points between D0 and D4 of differentiation. Data represent averages from 3-6 replicates and are presented as means +/- standard deviations (SD). The level of expression in scrambled day 0 samples were set to a value of 1 for genes *Ptn* and *Thbs2*, and other values are relative to that sample value. *, P≤0.05; **, P≤0.001; ***, P≤ 0.0001 (versus scrambled by Student's t test). (C) Reduced Prmt5 expression detected by Western blotting at day 0, day 1, and day 2 of 3T3-

L1 differentiation with *Prmt5* siRNA oligo3 (means±sd.; n=3/group); (right) quantification of protein expression. P-value calculated in relation to scrambled siRNA samples (*, $P \leq 0.05$; **, $P \leq 0.001$; ***, $P \leq 0.0001$) by Student's t-test.

A

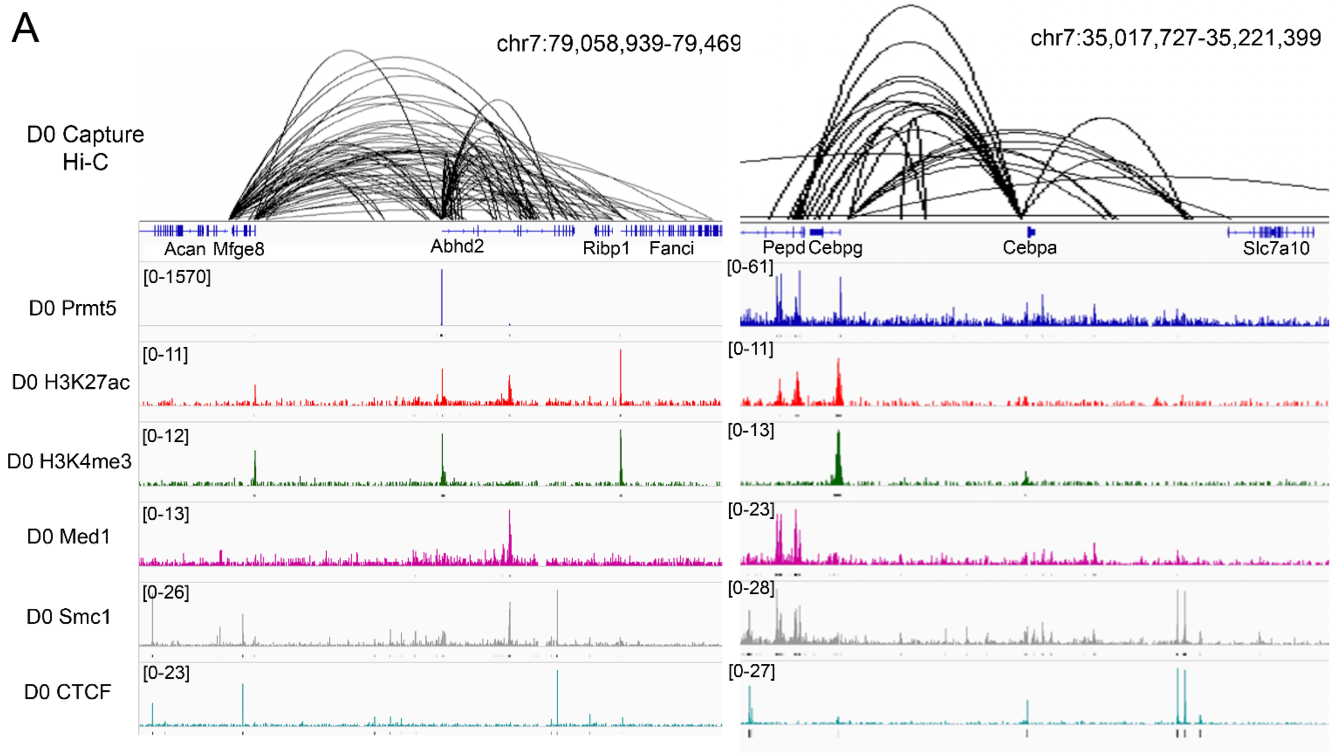


B

| Motif | Transcription Factor | p-value |
|---|---|---|
| 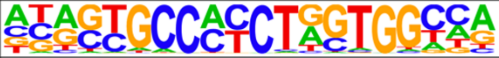 | CTCF(Zf) | 1.0E-175 |
| 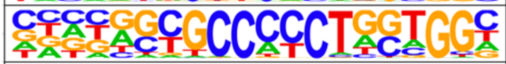 | BORIS(Zf) | 1.0E-126 |
| 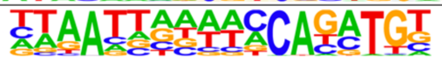 | Pitx1:Ebox | 1.0E-39 |
| 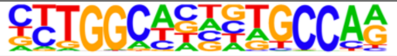 | NF1 | 1.0E-28 |

Supplemental Figure 4. Co-localization of Prmt5 with loop anchors and genome structure regulators. A) Genome Browser tracks showing Prmt5 ChIP-seq at day 0 of 3T3-L1 differentiation, along with H3K27ac, H3K4me3, Med1, Smc1, and Ctcf ChIP-Seqs with D0 Promoter Capture Hi-C displayed above for the *Abhd2* and *Cebpa* gene loci. (B) The HOMER motif discovery algorithm was performed on all Prmt5-bound high-confidence loop anchors and showed Ctcf and Boris as the top motifs present within peaks.
