## Supplementary material for "Protein arginine methyltransferase 5 (Prmt5) localizes to chromatin loop anchors and modulates expression of genes at TAD boundaries during early adipogenesis": Supp. Table 1

Supplemental Table 1.

**Table of read numbers for Prmt5 ChIP-seq**

| Sample Name | Biological Replicate | Mapped Reads |
| --- | --- | --- |
| D0 Input n1 | 1 | 58,533,683 |
| D0 Prmt5 ChIP n1 | 1 | 59,548,328 |
| D1 Input n1 | 1 | 49,099,767 |
| D1 Prmt5 ChIP n1 | 1 | 49,997,004 |
| D2 Input n1 | 1 | 52,525,844 |
| D2 Prmt5 ChIP n1 | 1 | 50,846,060 |
| D0 Input n2 | 2 | 53,936,052 |
| D0 Prmt5 ChIP n2 | 2 | 46,797,602 |
| D1 Input n2 | 2 | 46,183,452 |
| D1 Prmt5 ChIP n2 | 2 | 43,779,473 |
| D2 Input n2 | 2 | 54,745,955 |
| D2 Prmt5 ChIP n2 | 2 | 53,109,954 |

**Correlation of ChIP-Seq replicates**

| Biological Replicate 1 | Biological Replicate 2 | Pearson Correlation |
| --- | --- | --- |
| D0 Prmt5 ChIP n1 | D0 Prmt5 ChIP n2 | 0.89 |
| D1 Prmt5 ChIP n1 | D1 Prmt5 ChIP n2 | 0.59 |
| D2 Prmt5 ChIP n1 | D2 Prmt5 ChIP n2 | 0.68 |
