## Supplementary material for "Protein arginine methyltransferase 5 (Prmt5) localizes to chromatin loop anchors and modulates expression of genes at TAD boundaries during early adipogenesis": Supp. Tables 4-5

Supplemental Table 4

**Table of read numbers**

| Sample Name | Biological Replicate | Mapped Reads |
| --- | --- | --- |
| D0_HiC_scrambled_n1 | 1 | 121,752,254 |
| D0_HiC_oligo3_n1 | 1 | 107,852,797 |
| D0_ArimaHiC_scrambled_n2 | 2 | 100,011,523 |
| D0_ArimaHiC_oligo3_n2 | 2 | 105,298,239 |

**Correlation of replicates**

| Biological Replicate 1 vs Biological Replicate 2 | Pearson Correlation |
| --- | --- |
| chr1 | 0.983834 |
| chr2 | 0.985881 |
| chr3 | 0.972389 |
| chr4 | 0.984165 |
| chr5 | 0.984858 |
| chr6 | 0.986065 |
| chr7 | 0.928034 |
| chr8 | 0.984436 |
| chr9 | 0.98539 |
| chr10 | 0.98543 |
| chr11 | 0.985435 |
| chr12 | 0.982728 |
| chr13 | 0.984589 |
| chr14 | 0.982506 |
| chr15 | 0.987096 |
| chr16 | 0.984334 |
| chr17 | 0.985739 |
| chr18 | 0.984545 |
| chr19 | 0.985842 |
| chrX | 0.97149 |

**Supplemental Table 5 – Primers used**

| **siRNA** | **Catalog #** | **siRNA sequence** |
| --- | --- | --- |
| Prmt5 Oligo 3 Dharmacon | D-042281-04 | GUCCGUGCCUGUCGGGAAA |
| Prmt5 Oligo 5 (Invitrogen Stealth) | Custom, Prmt5 2075 | CCTACAGCACAGAAGGTGTAGAACA |

| **Expression Primer** |  |  |
| --- | --- | --- |
| Prmt5 | Forward | TGGGATGGCTGAAGGTAAAG |
|  | Reverse | TGAGGCACAGTTTGAGATGC |
| Pparγ2 | Forward | ATGCTGTTATGGGTGAAACTCT |
|  | Reverse | GGTAATTTCTTGTGAAGTGCTCATAG |
| Cebpa | Forward | CAAGAAGTCGGTGGACAAGAA |
|  | Reverse | CGTTGCGTTGTTTGGCTTTA |
| Thbs2 | Forward | GTATGGAGGGAAGGACTGTGT |
|  | Reverse | ACTTGGCTCCAGGAAAACACG |
| Ptn | Forward | TGGAGCTGAGTGCAAGTACC |
|  | Reverse | CTTTGACTCCGCTTGAGGCTT |
| Pdgfra | Forward | TCGCCAAAGTGGAAGAGACC |
|  | Reverse | CACCAGGACGATGAGAGAGA |
| Eef1a | Forward | GGCTTCACTGCTCAGGTGATTATC |
|  | Reverse | ACACATGGGCTTGCCAGGGAC |
